# Novel Association of Lyme disease, Age, and Atopic Dermatitis

**DOI:** 10.1101/2022.01.27.476641

**Authors:** Brandon T. Lee, Sarah D. Galloway, Qingying Feng, Satu Strausz, Maia Shoham, Paige Hansen, Laughing Bear Torrez Dulgeroff, Grace Blacker, Ying Y. Yiu, Paul Mansfield, FinnGen, Atif Saleem, Eric Gars, Erin C. Sanders, Irving L. Weissman, Hanna M. Ollila, Michal Caspi Tal

## Abstract

*Borrelia burgdorferi* (*B. burgdorferi*) is a bacterial spirochete that can cause Lyme disease after infecting a susceptible host. Immune responses to the bacteria are highly variable and host specific. The murine substrain, C3H/HeJ, is a frequently utilized mouse model of Lyme disease. In this study, we sought to investigate the correlation of age with onset and severity of dermatitis, both in mice infected with *B. burgdorferi* as well as humans who have had a diagnosis of Lyme disease. Female C3H/HeJ mice aged 6-8 weeks, 1 year, or 2 years were infected intraperitoneally with 10^5^ *B. burgdorferi*. Dermatitis of the tail was evaluated by gross examination and histology. Additional female C3H/HeJ and C57BL/6J mice aged 5 weeks were injected intradermally with 10^5^ *B. burgdorferi* containing the luciferase reporter gene then analyzed under in vivo imaging. Human data via electronic health records of 342,499 Finnish individuals was tested and analyzed for associations between Lyme disease and atopic dermatitis. Dermatitis worsened over the course of untreated infection, with ulceration, hemorrhaging, flaking, hair loss, and dark lesions as well as spongiosis and acanthosis. These features of dermatitis were present in infected mice after 1 year of age. We further confirm the presence of *B. burgdorferi* in the tail through quantification of bioluminescence and immunohistochemistry of both C3H/HeJ and C57BL/6J mice. This relationship among Lyme disease, atopic dermatitis, and host age seen in the mouse model is consistent with a large pool (342,499) of human epidemiological data from Finland. We identified 5,248 individuals with Lyme disease and 17,233 with atopic dermatitis in FinnGen. Retrospective analysis shows Lyme disease is associated with atopic dermatitis (OR = 1.91 [1.68 -2.37], *P* < 2e^-16^). More visits due to Lyme disease complications (3 or more visits versus 1 visit) were associated with atopic dermatitis (OR = 2.19 [1.35-3.55], *P* = 0.0014) and risk of developing atopic dermatitis over time (HR=2.26 [1.54-3.95], *P* = 0.0017). Data from mice and humans reveal a novel relationship among Lyme disease, age, and atopic dermatitis. Through defined pathological scoring, we demonstrate the onset of murine atopic dermatitis with *B. burgdorferi* infection, which is further exacerbated by host age at time of infection. In humans, a diagnosis of Lyme disease in FinnGen was associated with atopic dermatitis and further research is warranted to establish causation.

## INTRODUCTION

Lyme disease (LD) is the most prevalent tick-borne illness caused by infection with the spirochete *Borrelia burgdorferi*.^1^ LD is highly endemic in woody regions of the Northern Hemisphere with increasing prevalence.^2^ Patients may present with several different rash manifestations, including the hallmark “bulls-eye”, or no rash at all.^3,4^ Progression to the early disseminated stage of LD is characterized by various pathologies of the integumentary, musculoskeletal and nervous systems.^2,5^ If left untreated, late disseminated LD may present with cutaneous manifestations such as acrodermatitis chronica atrophicans, which may show variable atrophy and discoloration of skin.^6^

C3H/HeJ mice are commonly used as a model for LD because they develop several clinical features similar to human LD upon infection with *B. burgdorferi*, including carditis and arthritis.^7^ Compared to similar strains, C3H/HeJ mice exhibit chronic disease persistence with a higher degree of bacterial dissemination.^8^ In comparison, other inbred strains such as C57BL/6 mice do not exhibit as severe of infection, with lower antibody titers and no arthritis across the age.^7^ Interestingly, over the course of our studies on untreated long-term infection models of Lyme disease we noticed dermatitis with flaky lesions on the tail skin of C3H/HeJ mice after a year of infection with *B. burgdorferi*. However, the tail is not typically an organ of research interest and the cause of this dermatitis was unknown. Mouse studies in melanocyte pigmentation and psoriasis noted the similarity of epidermal layers between mouse tail skin and human skin, indicating the promising utility of the mouse tail as a model for pathology of the human epidermis.^9,10^

Atopic dermatitis (AD) is a chronic, inflammatory skin disease with usual onset occurring during childhood.^11,12^ This condition is characterized by pruritic, flaky lesions of the skin, hyperactive T^H^2-mediated immune responses, and relapsing episodes of skin inflammation.^13^ Symptoms vary with severity dependent on several genetic and environmental factors.^14,15^ Late onset AD diagnosed in the later life course tends to branch out of senescence, as increased “inflammaging” and decreased terminal keratinocyte differentiation, among other reported measures, are more robust than in early onset AD that manifests in childhood.^16–18^ Skin biopsies from patients with AD typically show varying degrees of spongiosis, acanthosis, and a superficial perivascular inflammatory infiltrate.^19^ Spongiosis is characterized by intraepidermal and intercellular edema which may visualized as increased space between epidermal keratinocytes on Hematoxylin and Eosin (H&E) stained histologic sections.^19^ Acanthosis is described as epidermal thickening.^20^ The perivascular inflammatory infiltrate (PVI) in atopic dermatitis is typically found in the superficial dermis and comprised of lymphocytes and histiocytes.^21^ Mast cells (MC) also may infiltrate the dermal layers of patients with AD, similar to individuals with inflammatory allergic responses.^14,22^ Furthermore, studies indicate C3H/HeJ mice present with PVI in the dura mater and ears, but less is known in the tail.^23,24^

Through histological analysis of phenotypic tails from infected mice, we define a scoring method to classify severity of presumed AD based on levels of spongiosis, acanthosis, and mast cell recruitment to the skin co-occurring with gross pathologic features of the tail exterior in C3H/HeJ and C57BL/6J mice. These scores show a significant pattern in severity of Lyme disease-associated atopic dermatitis associated with age. By using luciferase-containing spirochetes, we further observe the presence of *B. burgdorferi* in the tail under in vivo imaging. Furthermore, human epidemiological data from FinnGen, a large nationwide network of Finnish biobanks, support the findings we observe in the mouse model, showing a significant association between diagnoses of LD and AD.

## METHODS

### Mice

Female C3H/HeJ mice were purchased from Jackson Laboratories (Bar Harbor, ME) at ages of 6 weeks, 8 weeks, 1 year, or 2 years (n = 35). Experimental mice were infected with 10^5^ x *B. burgdorferi* spirochetes at 100 microliters (*μ*L) from a 7-day culture via intraperitoneal needle injection. All infected mice (n = 14) were age-matched to female, uninfected C3H/HeJ control mice (n = 18) that received a vehicle control of 100*μ*L of sterile Phosphate Buffered Saline (PBS) via intraperitoneal needle injection. Four mice expired prior to the completion of the experiment referenced in Figure 2, including uninfected 2 year (n = 2), uninfected 1 year (n = 1), and infected 1 year (n = 1). Humane euthanasia occurred either 2 months or 24 months post-injection for all experimental and control mice (n = 31) by 5% isoflurane vaporizer with active scavenging. Animal studies were performed at the Stanford School of Medicine Association for Assessment and Accreditation of Laboratory Animal Care International (AAALAC) accredited Rodent Animal Facility (Palo Alto, CA). All procedures and care guidelines were approved by the Stanford University Administrative Panel on Laboratory Animal Care (Protocol #30109).

Female C3H/HeJ and C57BL/6J mice were purchased from Jackson Laboratories at 6 weeks at time of infection. Experimental mice infected with 10^5^ x *B. burgdorferi* spirochetes at 50 microliters (*μ*L) resuspended in 0.2% mouse serum PBS (to reduce nonspecific interactions) from a 7-day culture via intradermal needle injection. 100*μ*L of double sterile filtered D-luciferin dissolved in sterile PBS (277mg/kg) was injected intraperitoneally per mouse 15 minutes prior to imaging for 1 minute exposure on In Vivo Imaging Spectrum Bioluminescent and Fluorescent Imaging Systems (IVIS) (Perkin Elmer) to confirm infection. Animal studies were also performed at the Department of Comparative (DCM) at the Massachusetts Institute of Technology (MIT) (Cambridge, MA). All procedures and care guidelines were approved by the MIT Committee on Animal Care (CAC) (Protocol#1221-087-24).

#### Borrelia burgdorferi

B31A3-GFP (Green Fluorescent Protein) *B. burgdorferi* were gifted by Dr. Jayakumar Rajadas of Stanford University in 10^7^ cells/mL aliquots and stored at -80º Celsius (C).^25^ ML23 containing a plasmid containing the firefly *luc* “luciferase” gene were generously provided by Dr Jenny Hyde of Texas A&M in 10^6^ cells/mL aliquots and stored at -80º Celsius.^26^ The spirochetes were thawed and cultured in 50 milliliter (mL) conical tubes (Corning) containing 50 mL Barbour-Stonner-Kelly (BSK-II) media complete with 6% rabbit serum (Millipore Sigma). Cultures were incubated at 37ºC in 5% carbon dioxide for 7 days. On the same day as *in vivo* infection, bacterial concentration was determined by Becton Dickinson (BD) LSRFortessa Flow Cytometer and cultures were adjusted to desired working concentration using PBS.

### Histology

Whole mouse tails were collected at sacrifice and fixed with 4% paraformaldehyde (PFA) for 24 hours before they were transferred to 70% ethanol. All subsequent tissue processing was outsourced to Histowiz (Brooklyn, NY) or to the MIT Histology Core Facility (Cambridge, MA). Tails were cut to show transverse cross sections of the most cranial, caudal, or medial region. Tissue was then stained with either H&E, anti-*Borrelia*, or Cluster of Differentiation 117 (CD117) antibody, a mast cell marker. All investigators and consulting pathologists were blinded for initial analyses to facilitate unbiased qualitative observations. Pathologists performed subsequent unblinded analysis of tail phenotypes utilizing recommendations set by the International Harmonization of Nomenclature and Diagnostic Criteria for Lesions in Rats and Mice (inHAND) Criteria.^27^

### Pathology Scoring & Quantification

Gross examination of the tails and microscopic analysis (Leica M205 FA) was performed to classify atopic dermatitis severity through a defined external scoring system from 0 to 5 based on increasing severity, with 5 being the most severe. Gross pathology and histopathology scoring schemes for murine skin models have been adapted from studies examining autoimmune encephalomyelitis and atopic dermatitis, assessing various phenotypes such as acanthosis and hyperkeratosis.^28,29^ Two clinical pathologists (A.S. and E.G) provided their expert guidance in support of these internal and external scores we describe here. Each progressively higher score includes pathologies of the previous tier. The parameters of this scale were gathered from previous studies of mouse phenotypes and initial analyses were performed blind. See 6 external categories below (Table 1):

**Table 1.** External Pathology Scoring.

| Tail External Pathology Scoring & Quantification |  |
| --- | --- |
| Score | Description |
| 0 | Normal |
| 1 | Areas of disorganized hair growth |
| 2 | Hair loss and dark lesions |
| 3 | Flaking with complete hair loss and dark lesions |
| 4 | Ulceration, hemorrhaging with flaking, hair loss, and dark lesions |
| 5 | Damaged distal tip of tail with ulceration, hemorrhaging, flaking, hair loss, and dark lesions |

Histology of the tails, as previously described, was used to classify severity of clinical features of atopic dermatitis through an internal scoring system. Spongiosis was scored in a binary fashion and acanthosis of the epidermis was classified below (Table 2):

**Table 2.**
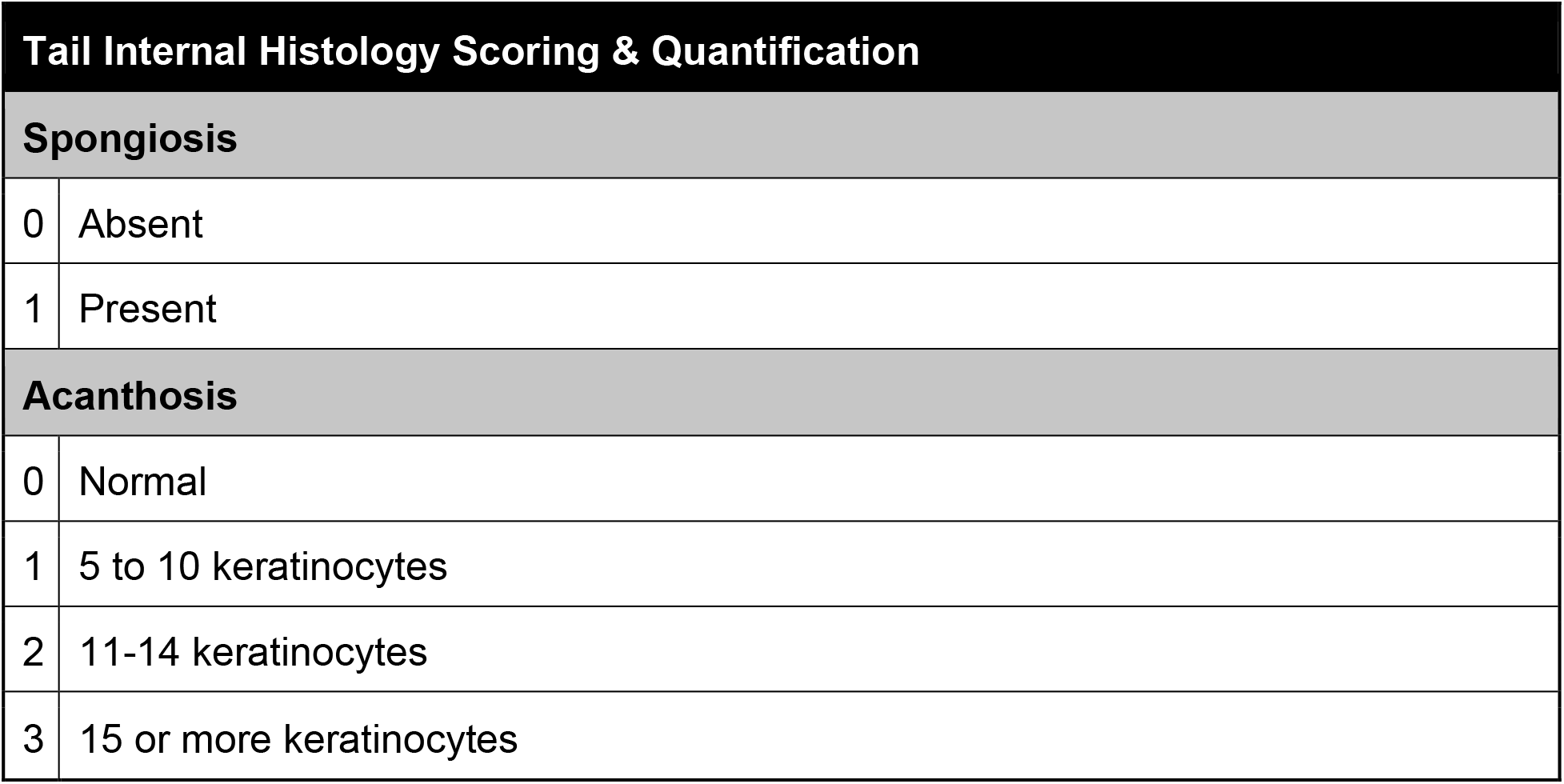
Internal Pathology Scoring.

| Tail Internal Histology Scoring & Quantification |  |
| --- | --- |
| Spongiosis |  |
| 0 | Absent |
| 1 | Present |
| Acanthosis |  |
| 0 | Normal |
| 1 | 5 to 10 keratinocytes |
| 2 | 11-14 keratinocytes |
| 3 | 15 or more keratinocytes |

Histologic scoring utilized the cranial cross section of each tail to maintain consistency across samples. Initial analyses were performed blinded with pathologist consultation, and identified variable degrees of spongiosis and acanthosis. There was no indication by their analysis of perivascular infiltrate in any of these samples. For each mouse, internal scores were summated and graphed against their respective external scores. Histology was conducted digitally, with files available upon request.

### Bioluminescence

At each subsequent time point from day 0 to 187, mice infected with ML23, the luciferase-containing strain of *B. burgdorferi*, were injected with 100 μL sterile filtered D-luciferin (GoldBio) reconstituted in sterile PBS at 277 mg/kg that was injected intraperitoneally. The hair on the backs of the mice was removed using an electric shaver, then the addition of Nair for 30 seconds to 1 minute. Gauze wipes were used to clean the backs in the following order: soaked in 70% ethanol, soaked distilled water, and a dry gauze. After 15 minutes, mice were arranged in IVIS for an image at an exposure time of 1 minute. Afterwards, total flux was quantified via Living Image Software and analyzed via R (v4.1.1) with the following packages: ggpubr, ggplot2, RColorBrewer.

### FinnGen

FinnGen is a large-scale study aiming to genotype 500,000 Finnish participants recruited from hospital samples as well as prospective and retrospective epidemiological and disease-based cohorts. This data is combined with longitudinal registries that record phenotypes and health events over the entire lifespan including the National Hospital Discharge Registry (inpatient and outpatient), Causes of Death Registry, the National Infectious Diseases Registry, Cancer Registry, Primary Health Care Registry (outpatient) and Drug Reimbursement Registry. This study used data from FinnGen Data Freeze 8, which includes 342,499 individuals.

We used data from the hospital inpatient, outpatient, primary outpatient, and drug reimbursement registries with International Classification of Diseases (ICD) codes for LD (ICD-10: A69.2, ICD-9: 1048A) and AD (ICD-10: L20, ICD-9: 6918 (6918X excluded), ICD-8: 691). We also retrieved information of sex, age at diagnosis, current age, and cohort. The nature of the registry information does not contain information of serology or chart data and we need to rely on the diagnoses made by the clinician.

Patients and control subjects in FinnGen provided informed consent for biobank research, based on the Finnish Biobank Act. Alternatively, separate research cohorts, collected prior the Finnish Biobank Act came into effect (in September 2013) and start of FinnGen (August 2017), were collected based on study-specific consents and later transferred to the Finnish biobanks after approval by Fimea (Finnish Medicines Agency), the National Supervisory Authority for Welfare and Health. Recruitment protocols followed the biobank protocols approved by Fimea. The Coordinating Ethics Committee of the Hospital District of Helsinki and Uusimaa (HUS) statement number for the FinnGen study is Nr HUS/990/2017.

The FinnGen study is approved by Finnish Institute for Health and Welfare (permit numbers: THL/2031/6.02.00/2017, THL/1101/5.05.00/2017, THL/341/6.02.00/2018, THL/2222/6.02.00/2018, THL/283/6.02.00/2019, THL/1721/5.05.00/2019 and THL/1524/5.05.00/2020), Digital and population data service agency (permit numbers: VRK43431/2017-3, VRK/6909/2018-3, VRK/4415/2019-3), the Social Insurance Institution (permit numbers: KELA 58/522/2017, KELA 131/522/2018, KELA 70/522/2019, KELA 98/522/2019, KELA 134/522/2019, KELA 138/522/2019, KELA 2/522/2020, KELA 16/522/2020), Findata permit numbers THL/2364/14.02/2020, THL/4055/14.06.00/2020,,THL/3433/14.06.00/2020, THL/4432/14.06/2020, THL/5189/14.06/2020, THL/5894/14.06.00/2020, THL/6619/14.06.00/2020, THL/209/14.06.00/2021, THL/688/14.06.00/2021, THL/1284/14.06.00/2021, THL/1965/14.06.00/2021, THL/5546/14.02.00/2020, THL/2658/14.06.00/2021, THL/4235/14.06.00/2021 and Statistics Finland (permit numbers: TK-53-1041-17 and TK/143/07.03.00/2020 (earlier TK-53-90-20) TK/1735/07.03.00/2021).

The Biobank Access Decisions for FinnGen samples and data utilized in FinnGen Data Freeze 8 include: THL Biobank BB2017_55, BB2017_111, BB2018_19, BB_2018_34, BB_2018_67, BB2018_71, BB2019_7, BB2019_8, BB2019_26, BB2020_1, Finnish Red Cross Blood Service Biobank 7.12.2017, Helsinki Biobank HUS/359/2017, Auria Biobank AB17-5154 and amendment #1 (August 17 2020), AB20-5926 and amendment #1 (April 23 2020), Biobank Borealis of Northern Finland_2017_1013, Biobank of Eastern Finland 1186/2018 and amendment 22 § /2020, Finnish Clinical Biobank Tampere MH0004 and amendments (21.02.2020 & 06.10.2020), Central Finland Biobank 1-2017, and Terveystalo Biobank STB 2018001.

### Data Analysis

We tested associations between LD and AD using logistic regression analysis utilizing the FinnGen data. The models were adjusted for age, sex and cohort. We calculated Cox proportional hazard model with age as the timescale using sex and cohort as covariates, and tested the model assumptions by cox.zph function. The total flux (photons per second, p/s) of each mouse is gated using the software LivingImage across different time points. We gated the regions of interest of the tails, establishing the minimum and maximum values of total flux detected by the IVIS for the entire bioluminscence study. In addition, we tested survival with Kaplan–Meier estimator using a non-parametric log-rank test. For the genetics-based study in FinnGen, we performed analyses in R version 4.1.0 using packages survminer and survival, and visualizing the results with survminer. A *P* value less than 0.05 was considered statistically significant.

For mouse models, GraphPad Prism v9.1.0 was used to develop and analyze graphs, utilizing two-way ANOVA and by Šidák’s multiple comparisons correction test for significance of *P* < 0.05.

## RESULTS

### 24-month-long chronic infection initiated in young mice leads to severe dermatitis of the tail

In our ongoing studies using a murine model of LD, we observed that many mice which had been infected with *B. burgdorferi*, and never treated with antibiotics, developed dermatitis. This dermatitis ranged in severity, with some aged animals developing dermatitis severe enough to warrant compassionate euthanasia. In order to determine the nature and cause of the dermatitis, and how it may relate to age, we infected 8-week-old female C3H/HeJ mice and closely examined them for 24 months, compared to uninfected age-matched control mice (Figure 1). Visualization of the mouse tail by light microscopy showed severe dermatitis in the infected mice. External features of the phenotypic tails included hair loss, skin flaking, hyperpigmentation, hemorrhaging, and ulceration.

**Figure 1.**
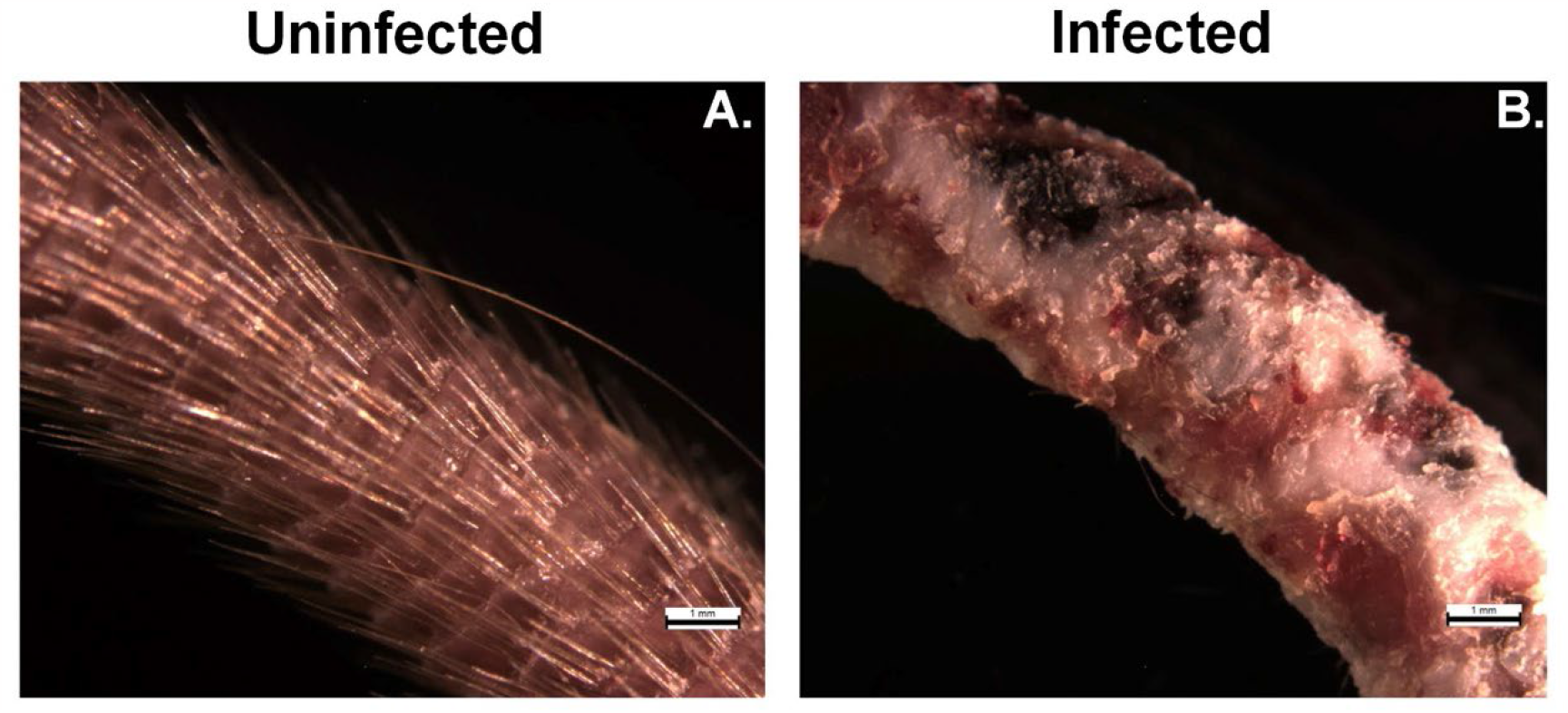
Chronic *B. burgdorferi* infection results in severe tail pathology. Uninfected female C3H/HeJ control mice (n = 4) (A) were compared to age-matched mice infected with B. burgdorferi at 8 weeks of age and sustained a chronic infection for 24 months (n = 3) (B). Upon euthanasia, tails were analyzed by light microscopy and evaluated for external pathology. Scale bars are equivalent to 1 mm.

**Figure 2.**
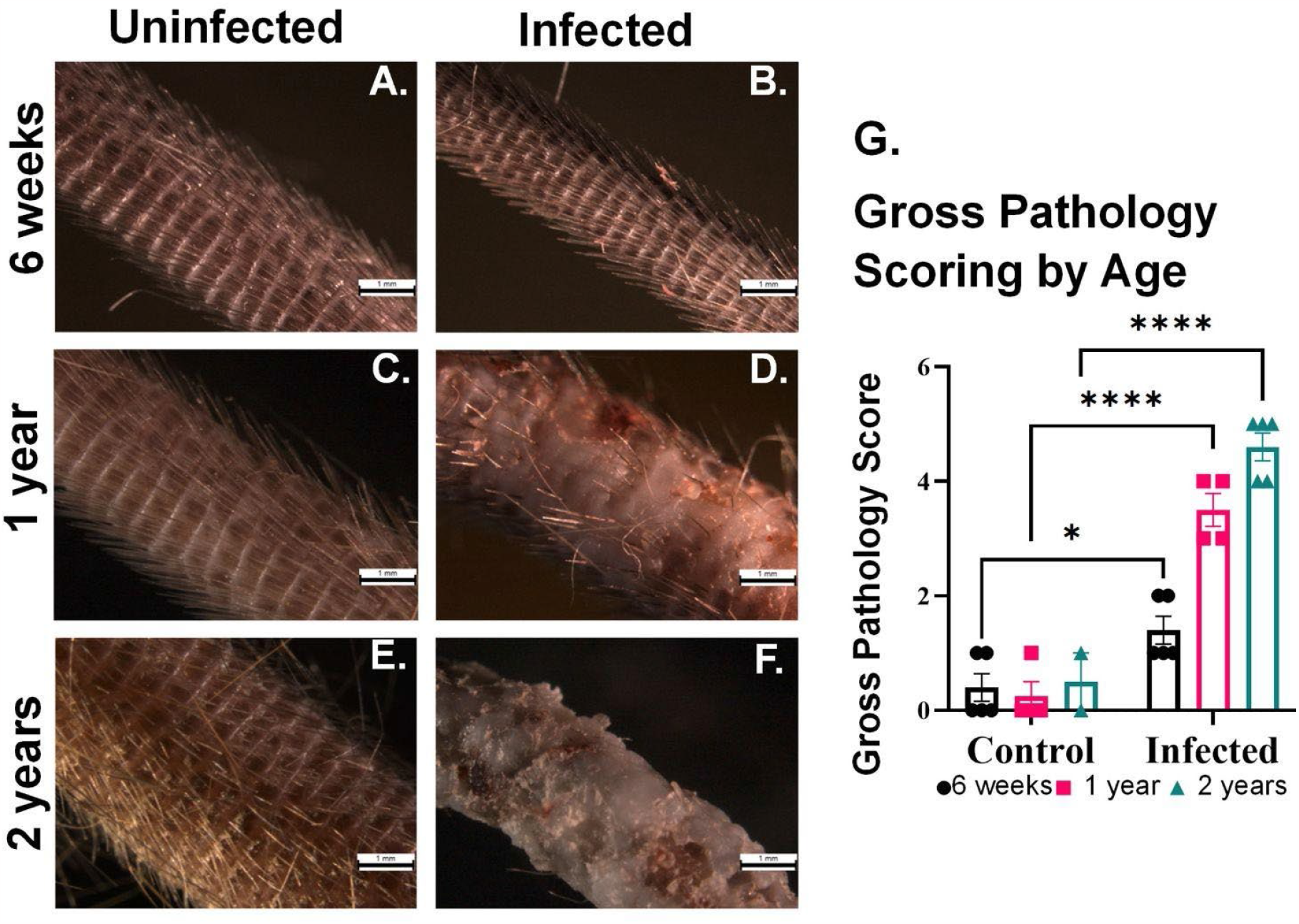
Gross tail pathology in mice with acute Lyme disease significantly increases with age. Uninfected female C3H/HeJ control mice aged 6 weeks (n = 5) (A), 1 year (n = 4) (C), or 2 years (n = 2) (E) were compared to mice infected with *B. burgdorferi* at equivalent ages of 6 weeks (n = 5) (B), 1 year (n = 4) (D), or 2 years (n = 5) (F). Images of tails were taken 2 months post-infection and severity of dermal pathology was evaluated and scored. Assigned scores were then analyzed using two-way ANOVA followed by Šidák’s multiple comparisons test (*P < 0.05, ****p < 0.0005) (G). Scale bars are equivalent to 1mm.

### Gross pathology scores of affected skin are significantly higher in aged mice with acute Lyme disease

Aging is known to cause immune system dysfunction and increased susceptibility to disease and infection.^30^ To further investigate the impact of age on the onset and severity of dermatitis symptoms in the tails of mice with LD, we set up an experiment of acute Lyme disease in female C3H/HeJ mice of various ages at the time of infection: 6 weeks, 1 year, and 2 years (Figure 2B, 2D, and 2F). Tails of infected mice were directly compared with age-matched uninfected controls: 6 weeks, 1 year, 2 years (Figure 2A, 2C, 2E). Two months post-infection, the mice were humanely euthanized and whole tails were evaluated by light microscopy (Figure 2). Mice infected at 6 weeks of age (Figure 2B) show a minimal difference in tail condition compared to age-matched, uninfected mice (Figure 2A). Mice infected at 1 year of age present with disorganized hair growth, varying degrees of hair loss, skin flaking, dark lesions and hemorrhaging (Figure 2D). Age-matched, 1-year-old uninfected controls show no pathology (Figure 2C). Mice infected at 2 years of age developed severe dermatitis with complete hair loss, skin flaking, dark lesions, hemorrhaging and damaged or missing distal tail tip (Figure 2F). Tails of 2-year-old age-matched, uninfected mice showed disorganized hair growth consistent with natural aging phenotypes of C3H/HeJ mice (Figure 2E). Surprisingly, the severe pathology of the acutely (2 month) infected 2-year-old mice (Figure 2F) is grossly indistinguishable from the chronically (24 month) infected mice (Figure 1B).

External pathology was quantified for all cohorts and analyzed using two-way ANOVA followed by Šidák’s multiple comparisons test (*P* < 0.005). Mice acutely infected at 6 weeks of age had minimal statistically significant differences in tail pathology to their uninfected controls (*P* <0.05, LS mean difference = 1.00, 95% CI [0.08154, 1.918]). Substantially greater statistical difference is seen in 1- and 2-year-old mice compared to their respective age-matched uninfected controls (*P* <0.0005, LS mean = 3.25, 95% CI [2.223, 4.277] and *P* <0.0005, LS mean = 4.10, 95% CI 3.885, 5.315]) (Figure 2G). Tail pathology external scoring and quantification parameters are detailed in the methods. Score values for tail pathologies in individual mice are shown in Supplementary Table 1.

### Features of atopic dermatitis found in histopathology of affected skin from mice acutely infected with B. burgdorferi show increased severity with advanced age

Human atopic dermatitis is commonly a clinical diagnosis and biopsies are only rarely performed in the evaluation.^11^ Similar to human histopathology, murine AD presents with varying levels of acanthosis, spongiosis, and perivascular inflammation.^20^ Another commonly observed feature of human atopic dermatitis, which is observed in the *B. burgdorferi-*infected mice of this study, is post-inflammatory hyperpigmentation of the skin, which is pigmentation that occurs after resolution of inflammatory skin eruptions.^31^ Gross pathology identified the development of age-dependent dermatitis on the tails of C3H/HeJ mice acutely infected (2 months) with *B. burgdorferi* in 3 different age groups (6 week, 1 year, 2 years). Tails from these mice were then processed for histology.

Transverse cross sections from the caudal, medial, and distal tail regions from these age-matched uninfected and infected mice were stained with H&E for evaluation (Supplementary Figure 1). Epidermal regions of the tail were closely evaluated for features of dermatitis (Figure 3). Mice infected with *B. burgdorferi* at 6 weeks of age showed lack of histopathologic findings (Figure 3B). Mice infected at 1 year of age developed acanthosis, spongiosis and mild thickening of the stratum corneum, as visualized by sections of the tail skin (Figure 3D). Mice infected at 2 years of age showed the most significant pathology with more severe acanthosis accompanied by elongated rete ridges, spongiosis and hyperkeratosis (Figure 3F). All mice infected at 1 and 2 years of age also showed loss of hair follicles and post-inflammatory hyperpigmentation consistent with observations from gross examination.

**Figure 3:**
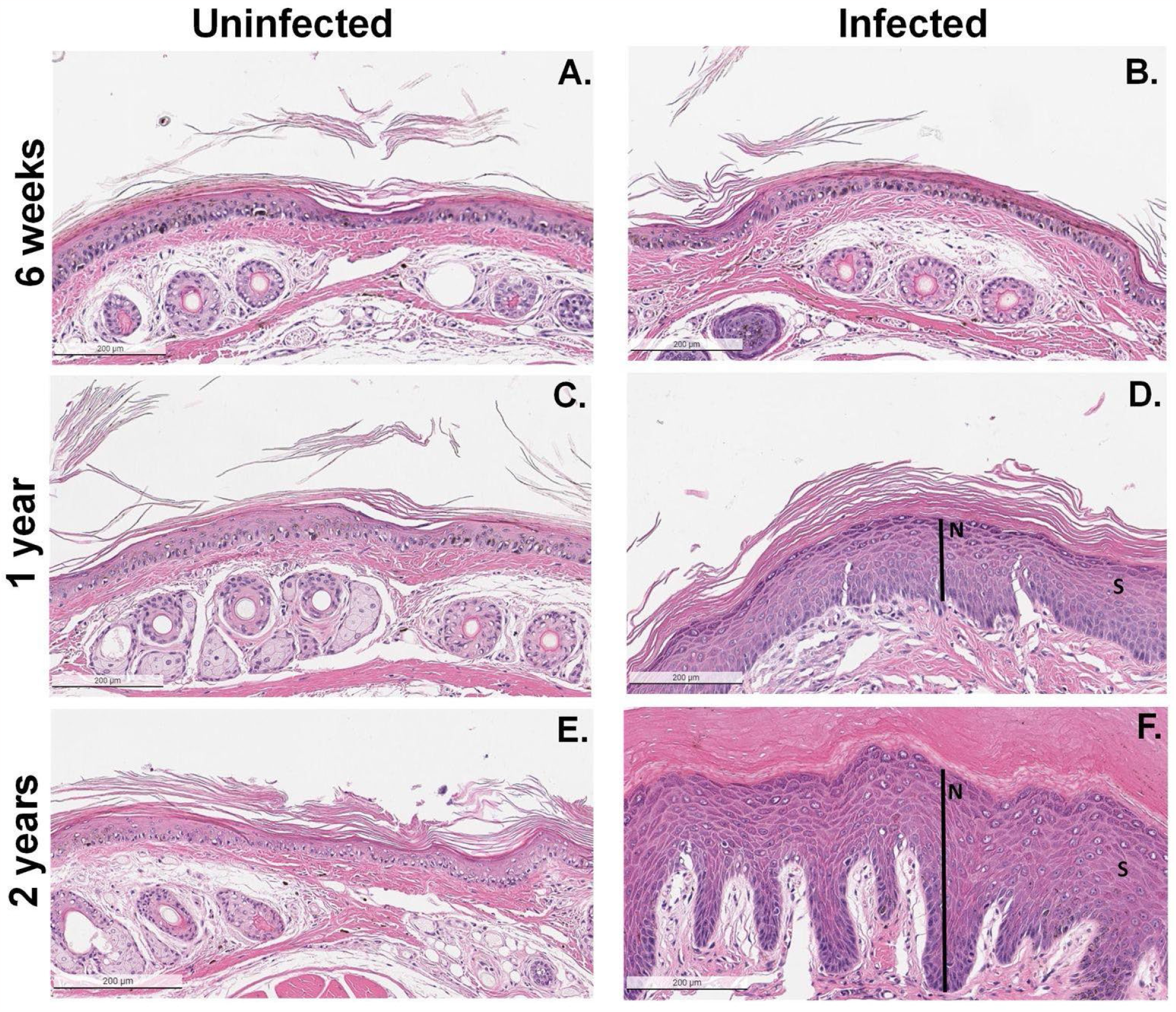
Tail histopathology of mice with acute *B. burgdorferi* infection reveal features of atopic dermatitis that worsen with age. Representative images of transverse cross sections of the mouse tail stained with H&E. Uninfected female C3H/HeJ control mice aged 6 weeks (n = 5) (A), 1 year (n = 4) (C), or 2 years (n = 2) (E) were compared to mice infected with B. burgdorferi at equivalent ages of 6 weeks (n = 5) (B), 1 year (n = 4) (D), or 2 years (n = 5) (F) Mice were humanely euthanized at 2 months post-infection and all tails were collected for histology. Presence of spongiosis is exemplified by an “S”, and acanthosis of the epidermis is indicated by a black line labeled “N”. All images are at 10X magnification, and scale bars are equivalent to 200μm.

The assigned scores for both external and internal pathology were then compared via heat map which revealed strong associations (Supplementary Figure 2). Individual mice with severe external features of AD additionally had congruent histopathologic findings. Likewise, uninfected mice had low scores for both external and internal AD pathology.

### Mast cells localize to atopic dermatitis lesions consistent with age-dependent severity in mice with acute Lyme disease

Mast cells (MC) are key mediators of the T^H^2 immune response and are classically involved with allergy and IgE-mediated inflammation.^32^ Several studies have implicated MC involvement in AD pathogenesis, including IgE-mediated sensitization to environmental allergens as a typical feature of human AD.^22^ Further, *B. burgdorferi* spirochetes are capable of inducing mast cell degranulation through interactions with proteins found on the spirochete membrane.^33,34^ One such defined interaction is through lipidated outer surface protein C lipoprotein, found on B. burgdorferi, which has been shown to induce degranulation of primary murine mast cells, but was nonessential to mast cell activation by intact *B. burgdorferi* spirochetes. We therefore investigated if the age-dependent pattern of AD severity seen in C3H/HeJ mice with LD may be correlated with MC localization to these lesions.

Tail sections from mice infected with *B. burgdorferi* at ages of 6 weeks, 1 year, and 2 years along with their aged matched uninfected controls were stained antibodies against CD117, a receptor tyrosine kinase and mast cell marker (Figure 4A-F). During analysis, mast cells were classified by cellular shape and CD117 stain intensity. All positively identified MC within the dermis of a single tail section were quantified for all cohorts and analyzed using two-way ANOVA followed by Šidák’s multiple comparisons test (P < 0.005).

**Figure 4:**
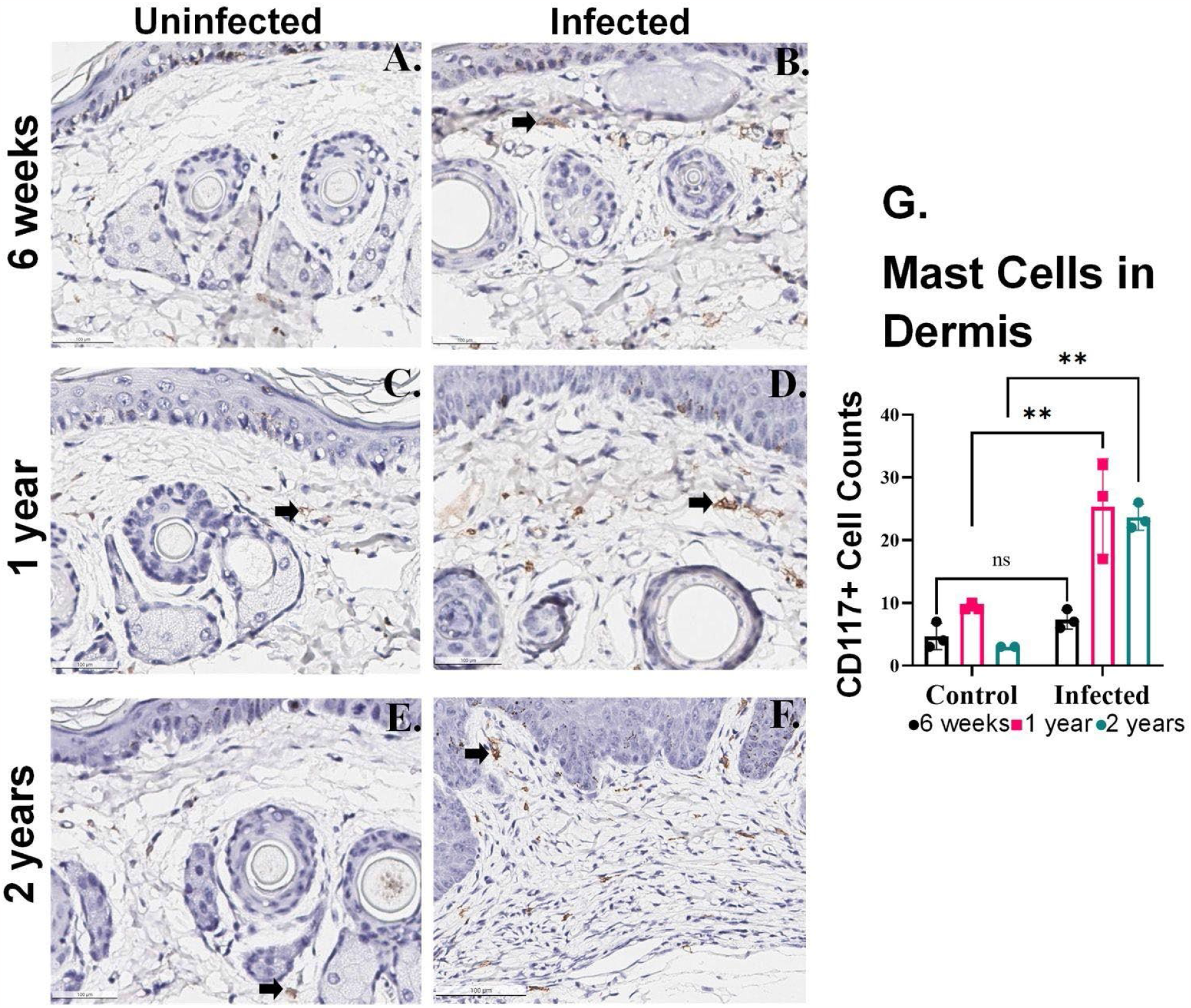
Mast cells localize to atopic dermatitis lesions consistent with age-dependent severity in mice with acute Lyme disease. Representative images of transverse cross sections of the mouse tail stained for CD117 (brown) with a hematoxylin counterstain (purple). Uninfected female C3H/HeJ control mice aged 6 weeks (n = 3) (A), 1 year (n = 3) (C), or 2 years (n = 2) (E) were compared to mice infected with *B. burgdorferi* at equivalent ages of 6 weeks (n = 3) (B), 1 year (n = 3) (D), or 2 years (n = 3) (F). Mice were humanely euthanized at 2 months post-infection and a select number of tails were collected for histology to determine mast cell prevalence by CD117 staining. Black arrows indicate a representative mast cell in each image. The total mast cell count is shown by age and condition using two-way ANOVA followed by Šidák’s multiple comparisons test (ns p > 0.05, **P < 0.005) (G). Scale bars are equivalent to 100μm.

Interestingly, both acutely infected 1- and 2-year old mice groups had substantial MC localization to the AD lesions (Figure 4G). Mice infected at 1 year of age (*P* = 0.005, LS mean = 16, 95% CI 6.715, 25.29]) or 2 years of age (*P* = 0.0002, LS mean = 20.72, 95% CI [10.17, 31.28]) had significantly more MCs in the dermal layer of the tail than their age-matched, uninfected controls. In contrast, mice infected at 6 weeks of age had no significant difference in MC count compared to their respective uninfected controls (*P* = 0.74, LS mean = 2.667, 95% CI [-6.619, 11.95]).

### Lyme disease associates with atopic dermatitis in humans

Recent large scale cohorts have provided means to explore epidemiological correlates leveraging on electronic health records. Using electronic health records comprising 342,499 individuals in FinnGen we identified 5,248 individuals with LD; 444 individuals from hospital inpatient, 1,317 individuals from hospital outpatient and 3,487 individuals from primary outpatient registries. We identified 17,233 individuals with AD; 3,063 individuals from hospital inpatient, 7,841 individuals from hospital outpatient, 5,796 individuals from primary outpatient and 533 individuals from drug reimbursement registries with 298 individuals with both LD and AD, and the majority of individuals being disease free for LD or AD (n = 320,316) (Figure 5A).

**Figure 5:**
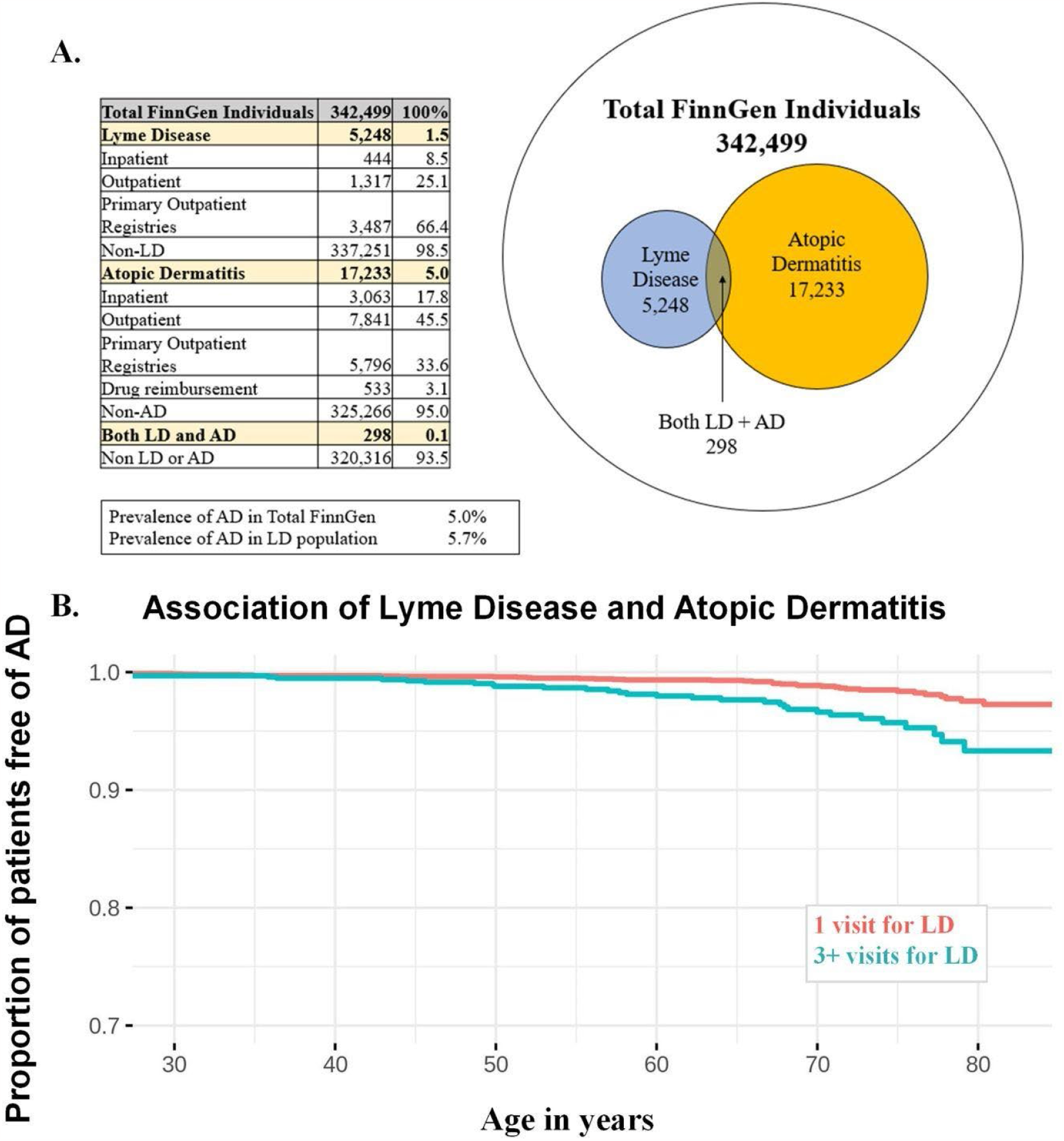
Association of Lyme disease (LD) with atopic dermatitis (AD) in FinnGen. (A) Prevalence of LD and AD in FinnGen. (B) Kaplan-Meier survival analysis in incident AD in individuals with 1 diagnosis for LD (blue), or at least 3 LD diagnoses (red). All individuals (N = 75) have at least one LD diagnosis prior to onset of AD (log rank P = 4.0e-5).

Overall, we identified 298 prevalent cases that had diagnosis for both AD and LD, and 85 incident cases that had AD diagnosis after LD diagnosis. AD was more frequent in individuals with LD (5.7% in individuals with LD vs. 5.0% in individuals without LD) (Figure 5A). Our study shows an epidemiological association between Lyme disease and AD with higher odds of AD in patients with Lyme disease.

(OR = 1.91 [1.68 -2.37], *P* < 2e-16). We estimated the severity of LD infection by dividing the number of diagnoses into two categories; 3 or more LD diagnoses vs. 1 LD diagnosis. We excluded individuals who had AD diagnosis prior to LD diagnosis (N remaining = 4,373 individuals with LD, 75 participants with LD and AD diagnosis). First, we calculated logistic regression model and discovered that LD with more than 3 diagnoses was significantly associated with increased risk for AD (OR=2.19[1.35-3.55], *P*=0.0014). In addition, we estimated temporal effect by Cox-proportional hazard model using age as the timescale showing corresponding result (HR=2.26[1.54-3.95], *P*=0.0017). Finally, we calculated Kaplan Meier estimates demonstrating that survival probability differed significantly between groups 3 or more LD diagnoses vs. 1 LD diagnosis. Consequently, we show that patients in the 1 LD diagnosis group had a higher AD free survival percentage compared to individuals in the 3 or more LD group (P = 4.0e-5) (Figure 5B).

### *In vivo* tracking of *B. burgdorferi* localizes into the tail

While other bacteria have been found to establish infection of the tail, this trophism has not been previously reported in *B. burgdorferi*.^35,36^ To ask whether *B. burgdorferi* were co-localized at the sites of atopic dermatitis, bacteria were imaged using an In Vivo Imaging System (IVIS) to visualize and quantify the bacterial load through the mice. The *B. burgdorferi* strain ML23-pBBE22*luc* (ML23) was used for infection, as it contains a luciferase plasmid reporter. This engineered ML23-LUC strain emits bioluminescence in an ATP-dependent reaction when live bacteria are exposed to the luciferin substrate and imaged under the IVIS machine with adequate exposure time. Weekly imaging was conducted to track the dissemination of *B. burgdorferi*, which showed the location of *B. burgdorferi* in infected mice.

Importantly, both C3H and B6 female mice had measurable luciferase signals in the tail three weeks after initiation of infection (Figure 6). The total flux (p/s) from the tails of C3H mice were significantly higher than in those of the B6 mice (Two-Way Anova, F-value = 15.28, p = 0.000111). (Supplemental Figures 3–6).

**Figure 6.**
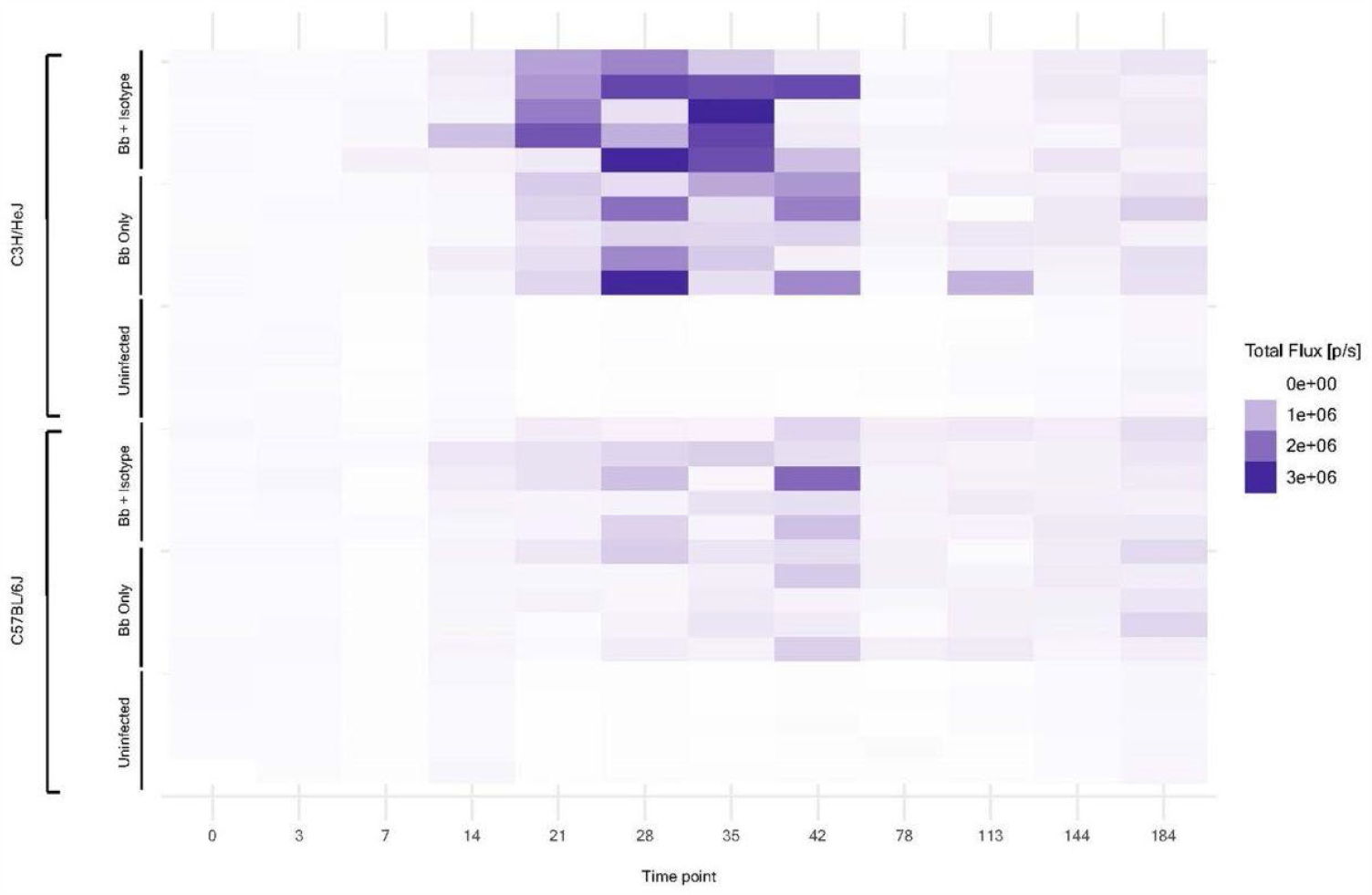
Quantified bacterial load of mice infected with luciferase-containing *B. burgdorferi* in the tails. C57BL/6J and C3H/HeJ mice aged at x time of infection were infected with ML23 *B. burgdorferi* intra-dermally and monitored over 187 days. Bioluminescence was measured by total flux through MS, which detects signal from luciferase-activity stimulated by a 15 minute incubation of D-luciferin dissolved in sterile PBS, injected intraperitoneally al 277mg/kg per mouse.

## DISCUSSION

Classic phenotypes of Lyme disease in humans after acute *B. burgdorferi* infection include skin manifestations of various clinical phenotypes such as the erythema migrans, but this varies significantly across humans. New onset of chronic dermatologic conditions after LD have not been well investigated. The C3H/HeJ mouse model is frequently used in LD research because it presents with both arthritis and carditis.

However, we observed that female C3H/HeJ mice chronically infected with *B. burgdorferi* often developed severe dermatitis of the tail. Since the mouse tail skin shares more similarity with human skin as compared to areas of skin with dense fur, we focused on this tissue to investigate this dermatitis further in both C3H/HeJ and C57BL/6J mice. Due to the absence of noticeable dermatitis in younger infected mice, we investigated whether dermatitis in older infected mice was primarily driven by the length of infection or the age of the host animal at the time of infection.

We developed a numeric scoring system to assess the external pathology and internal histology of murine skin. Higher scores were associated with more severe pathology. Both internal and external scoring increasingly worsened with age and we found a strong correlation between increased age and AD severity of the tail among *B. burgdorferi-*infected mice. Interestingly, internal and external scores of both acutely (2 months) infected 2-year-old mice, and mice who had been chronically (24 months) infected since 8 weeks old, both resulted in severe pathologic changes of hair loss, skin flaking, hyperpigmentation, and ulceration. The youngest mice (6 weeks old) did not develop features of AD within the 2-month observation period before euthanasia. While natural aging does impact hair growth and tissue quality in uninfected controls, the infection consistently and significantly amplified the extent of AD lesions on the tails of mice with LD. These lesions were also associated with increased mast cell infiltration to the tail dermis. Other signs noted were mild pruritus and ankle swelling that varied across cages and infected mice. We propose that utilizing similar gross and histopathological scoring systems will supplement long-term monitoring of mice infected with *B. burgdorferi* at the tail.

Given the dermatologic findings we observed in both acutely and chronically infected aged mice, we were curious to see if Lyme disease was also associated with atopic dermatitis in humans through analysis of clinically diagnosed LD and AD patients from FinnGen in 342,499 individuals. We found a significant association between LD and AD diagnoses that warrant further investigation into this relationship we report here. Among people with LD, there was a higher prevalence of AD than among the general population. Approximately 90% of individuals who are diagnosed with acute LD recover without complications, and therefore we further divided the individuals with LD in FinnGen into a subset who had one medical visit due to LD and a subset who had three or more visits. We observed an association with AD and the number of visits due to LD, which may reflect disease severity. Consequently, those individuals who have several healthcare visits due to LD may have a higher risk for AD than individuals with only one visit.

Increased age can have damaging effects on immune system regulation and host defenses, while also raising the risk for inflammatory conditions.^32^ Age-related defects in critical mechanisms such as production of pro-inflammatory cytokines (TNF-α and IFN-γ) and terminal differentiation of keratinocytes can induce differential phenotypes of the epidermis.^33^ Mast cells are a major effector cell type implicated in pathogenesis of human AD through the release of proinflammatory mediators and IgE-mediated sensitization of environmental allergens.^22^ Utilizing mouse models, mast cells have also been implicated in other age-related pathologic mechanisms, such as damage to lung endothelial tissue due to MC-mediated neutrophil migration.^37^ Existing literature exploring the role of MC in murine LD is limited; however, evaluation of gerbils infected with *B. burgdorferi* showed significant histopathologic changes, including MC infiltration of the extracellular matrix surrounding major organs.^38^ As LD is an illness primarily characterized by inflammation of the joints, infection with *B. burgdorferi* could contribute to mast cell dysregulation and possibly lead to a higher risk of developing inflammatory diseases such as AD.

Aged female C3H/HeJ mice with acute and chronic LD present with AD of the tail that is indistinguishable by gross evaluation and histology; a finding that was not observed in uninfected age-matched controls or in young mice. Importantly, *B. burgdorferi* is found at the site where AD later develops, and can be measured longitudinally using IVIS imaging. The degree of the tail-specific pathology is more concentrated in the tails post dissemination of *B. burgdorferi* throughout the mammalian host.^39^ This phenotype establishes a significant association between age of host with LD and onset of AD. We provide a numeric scale to quantify severity of AD, both internally and externally. This work also supports utilization of the C3H/HeJ mouse model in further study of Lyme-disease associated AD as compared to strains such as C57BL/6J mice that have a less pronounced response to LD overall. The C57BL/6J mice do not develop notable AD, but do develop dermatitis at other sites of the body, which may be related to their mild response to LD.^7^ The corroborating human data from FinnGen also suggests a novel relationship among age, Lyme disease, and atopic dermatitis through this epidemiological study and remains to be explored in future studies.

### LIMITATIONS

Symptoms of hair loss, ulceration, and tail tip disfiguration could be exacerbated by mouse self-mutilation induced from pruritus, or itch. Mice used for this study were not monitored for behavior, so frequency and severity of self-scratching or biting cannot be determined. When external tail pathology was observed, it was present in all mice of a single cage, suggesting that the pathology was not due to aggressive behaviors between individual mice.

The human disease association data from FinnGen may reflect more severe pathology since FinnGen is enriched for hospital level diagnoses. The nature of the registry information does not contain information of serology or chart data and thus analysis is dependent on the information encoded by their respective ICD codes for LD (ICD-10: A69.2, ICD-9: 1048A) and AD (ICD-10: L20, ICD-9: 6918 (6918X excluded), ICD-8: 691), based on the diagnoses input by the clinician into the medical record. Therefore, it cannot be said that these findings from our study indicate trends across individuals, rather broader populations of patients. The enrichment for hospital level diagnoses may result in larger effect sizes than what may be observed in more benign pathology or in outpatient only data. Our study shows higher odds of AD in patients with Lyme disease, however, while temporally related, our study does not directly show a causal association in humans.

Furthermore, the number of LD diagnoses have increased substantially during the last ten years. In the sample population found in FinnGen, the median age of patients diagnosed with AD was 32.9 years, and the median age of the LD population was 63.9 years. The recent shift in frequency of diagnoses of LD and the earlier age of diagnosis in AD may affect our power to examine any temporal associations between LD and AD that we conduct in this patient population. These associations are only suggestive and warrant further investigation at the basic and clinical science levels to validate this relationship we extrapolated from a small data set with some statistical significance that will be further explored with additional studies. Some of these limitations can be addressed in the future with larger data sets and with longer follow-up.

## DECLARATIONS

### ETHICS APPROVAL/CONSENT TO PARTICIPATE

Animal studies were performed at the Stanford School of Medicine Association for Assessment and Accreditation of Laboratory Animal Care International (AAALAC) accredited Rodent Animal Facility (Palo Alto, CA). All procedures and care guidelines were approved by the Stanford University Administrative Panel on Laboratory Animal Care (Protocol #30109).

## AVAILABILITY OF DATA & MATERIALS

All data and associated analyses conducted within this study will be made available by the corresponding author upon reasonable request.

## CONFLICT OF INTEREST

None of the authors have any conflicts of research regarding the research presented in this manuscript.

## FUNDING

Research reported in this publication was supported by the Fairbairn family foundation; the Younger family foundation; the Robert J. Kleberg, Jr., and Helen C. Kleberg Foundation; Bay Area Lyme Foundation; the Virginia and D. K. Ludwig Fund for Cancer Research; M.C.T. and Y.Y.Y. were supported by Stanford Immunology training grant 5T32AI007290, and M.C.T. was also supported by the NIH NRSA 1 F32 AI124558-01 award. S.D.G. was supported by the California Institute for Regenerative Medicine Bridges 2.0 Grant EDUC2-08397. L.B.T.D. was supported by a Stanford Diversifying Academia Recruiting Excellence Fellowship.

The FinnGen project is funded by two grants from Business Finland (HUS 4685/31/2016 and UH 4386/31/2016) and the following industry partners: AbbVie Inc., AstraZeneca UK Ltd, Biogen MA Inc., Bristol Myers Squibb (and Celgene Corporation & Celgene International II Sàrl), Genentech Inc., Merck Sharp & Dohme Corp, Pfizer Inc., GlaxoSmithKline Intellectual Property Development Ltd., Sanofi US Services Inc., Maze Therapeutics Inc., Janssen Biotech Inc, Novartis AG, and Boehringer Ingelheim. Following biobanks are acknowledged for delivering biobank samples to FinnGen: Auria Biobank (www.auria.fi/biopankki), THL Biobank (www.thl.fi/biobank), Helsinki Biobank (www.helsinginbiopankki.fi), Biobank Borealis of Northern Finland (https://www.ppshp.fi/Tutkimus-ja-opetus/Biopankki/Pages/Biobank-Borealis-briefly-in-English.aspx), Finnish Clinical Biobank Tampere (www.tays.fi/en-US/Research_and_development/Finnish_Clinical_Biobank_Tampere), Biobank of Eastern Finland (www.ita-suomenbiopankki.fi/en), Central Finland Biobank (www.ksshp.fi/fi-FI/Potilaalle/Biopankki), Finnish Red Cross Blood Service Biobank (www.veripalvelu.fi/verenluovutus/biopankkitoiminta) and Terveystalo Biobank (www.terveystalo.com/fi/Yritystietoa/Terveystalo-Biopankki/Biopankki/). All Finnish Biobanks are members of BBMRI.fi infrastructure (www.bbmri.fi). Finnish Biobank Cooperative -FINBB (https://finbb.fi/) is the coordinator of BBMRI-ERIC operations in Finland. The Finnish biobank data can be accessed through the Fingenious^®^ services (https://site.fingenious.fi/en/) managed by FINBB. The funders had no role in study design, data collection and analysis, decision to publish, or preparation of the manuscript.

## AUTHORS’ CONTRIBUTIONS

B.T.L. M.S., and Q.F. carried out infections and observations. B.T.L., L.B.T.D., P.M., P.H., and Y.Y.Y. conducted endpoint tissue procurement. S.G. prepared samples for histopathology and contributed to histopathology analysis. A.S. and E.G. conducted primary analysis of histopathology and murine diagnosis. The FinnGen Project was utilized for human epidemiological data that was studied and analyzed by S.S. and H.M.O. S.G. wrote the manuscript with B.T.L. Subsequent manuscript contributions and edits were made by M.C.T., E.S., S.S., H.M.O., P.H., G.B, and Q.F. B.T.L., Q.F., S.S., and H.M.O. created figures with contributions by M.C.T., S.G., G.B., E.S., and P.H.. M.C.T. and I.L.W. supervised all experiments and writing of the manuscript. All authors read and approved the final manuscript.

## ACKNOWLEDGEMENTS

The authors wish to thank members of the Weissman Lab at the Institute for Stem Cell Biology & Regenerative Medicine at Stanford University School of Medicine and Histowiz for their processing of our histological samples. The authors would also like to thank Caroline Atkinson and Sarah Elmiligy of the Preclinical Imaging Core at the Massachusetts Institute of Technology for their assistance in histology and in vivo imaging for subsequent experiments with luciferase. We want to acknowledge the participants and investigators of the FinnGen study.

## Notes

### Competing Interest Statement

The authors have declared no competing interest.

### Summary of Updates

Author contributions were modified to reflect contributions of the third author, who joined this study in the design and analysis of novel experiments that involve in vivo bioluminescence imaging of luciferase-containing B. burgdorferi. A larger emphasis on aging as an impactful factor in Lyme disease pathogenesis is placed in this revision than compared to the previous iteration. Each section has additional citations, and additional supplemental files are included to show the bioluminescence in mice.

