## Supplemental Figures for "Novel Association of Lyme disease, Age, and Atopic Dermatitis"

**A.**

| | 6 weeks | | 1 year | | 2 years | |
|---|---|---|---|---|---|---|
| | Control (n = 5) | Infected (n = 5) | Control (n = 4) | Infected (n = 4) | Control (n = 2) | Infected (n = 5) |
| **Disorganized hair growth** | 0/5 | 4/5 | 0/4 | 4/4 | 1/2 | 5/5 |
| **Hyperpigmentation** | 0/5 | 0/5 | 0/4 | 4/4 | 0/2 | 5/5 |
| **Hair loss** | 0/5 | 0/5 | 0/4 | 3/4 | 1/2 | 5/5 |
| **Flaking** | 0/5 | 0/5 | 0/4 | 4/4 | 0/2 | 5/5 |
| **Hemorrhage** | 0/5 | 0/5 | 0/4 | 2/4 | 0/2 | 4/5 |
| **Damaged tip of tail** | 0/5 | 0/5 | 0/4 | 0/4 | 0/2 | 3/5 |
| **Ulceration** | 0/5 | 1/5 | 0/4 | 3/4 | 0/2 | 5/5 |

**B.**

| | 6 weeks | | 1 year | | 2 years | |
|---|---|---|---|---|---|---|
| | Control (n = 5) | Infected (n = 5) | Control (n = 4) | Infected (n = 4) | Control (n = 2) | Infected (n = 5) |
| **Acanthosis** | 0/5 | 0/5 | 0/4 | 4/4 | 0/2 | 5/5 |
| **Flaking stratum corneum** | 0/5 | 0/5 | 0/4 | 4/4 | 1/2 | 5/5 |
| **Loss of dermal hair follicles** | 0/5 | 0/5 | 0/4 | 3/4 | 0/2 | 5/5 |
| **Spongiosis** | 0/5 | 0/5 | 0/4 | 3/4 | 0/2 | 5/5 |

**Supplementary Table 1: Pathologic scoring quantification of tails in mice with acute Lyme disease.** (A) External pathology. Each condition observed on the mouse tail and used in external scoring is reported as present or absent per individual mouse. Top row indicates age of mice at time of infection with B. burgdorferi or injection with vehicle control. (B) Internal Histopathology. Each condition observed in the mouse tail and used for internal scoring is reported as present or absent per individual mouse. Top row indicates age of mice at time of infection with B. burgdorferi or injection with vehicle control.

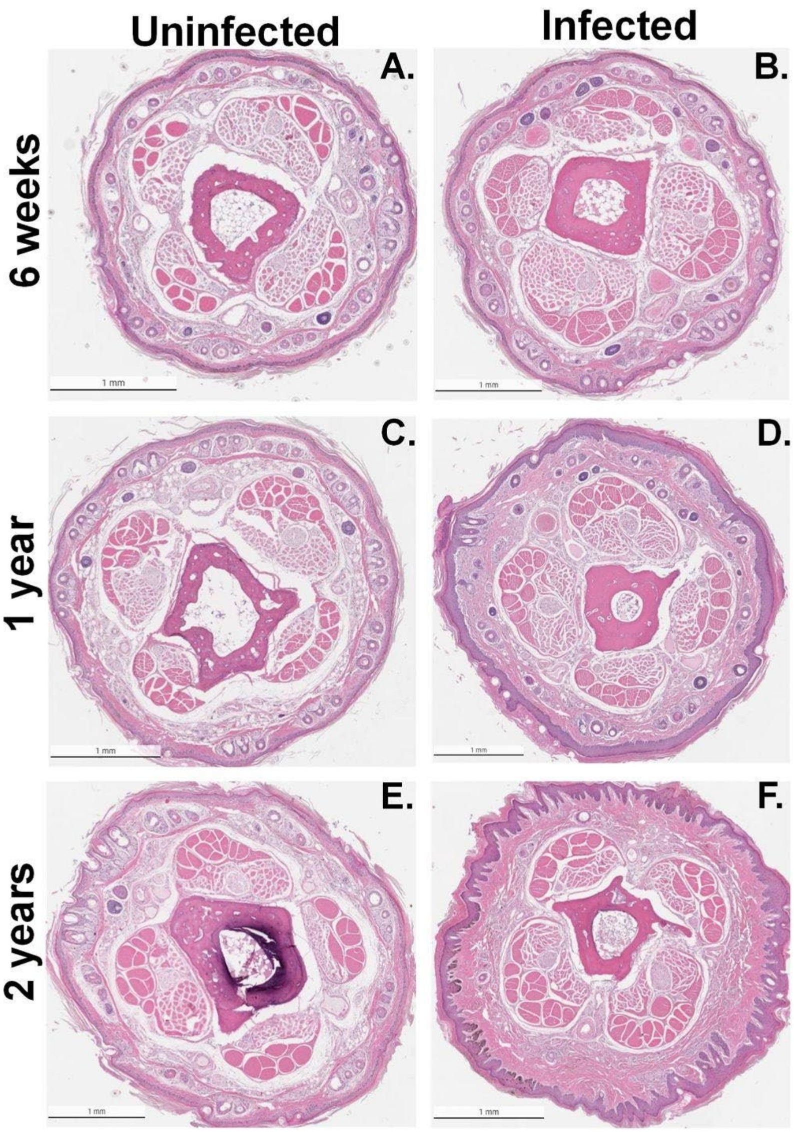


**Supplementary Figure 1: Histopathology of tail cross sections by age**. Representative images of entire tail cross sections from mice infected with *B. burgdorferi* at ages of 6 weeks (B), 1 year (D) or 2 years (F) that sustained an acute infection for 2 months. Respective age-matched, uninfected mice given vehicle controls are shown for comparison (A, C, E). Scale bars are equivalent to 1mm.

| ABBREVIATIONS | |
|---|---|
| AD | Atopic Dermatitis |
| ACA | Acrodermatitis Chronica Atrophicans |
| AAALAC | Association for Assessment and Accreditation of Laboratory Animal Care International |
| *B. burgdorferi* | *Borrelia burgdorferi* |
| BD | Becton Dickinson |
| BSK-H | Barbour-Stonner-Kelly with 4-(2-hydroxyethyl)-1-piperazineethanesulfonic acid (HEPES) buffer |
| C | Celsius |
| CD117 | Cluster of Differentiation 117 |
| CDC | Centers for Disease Control and Prevention |
| DEJ | Dermal-Epidermal Junction |
| DNCB | 2,4-dinitrochlorobenzene |
| GFP | Green Fluorescent Protein |
| H&E | Hematoxylin and Eosin |
| ICD | International Classification of Diseases |
| IL | Interleukin |
| inHAND | International Harmonization of Nomenclature and Diagnostic Criteria for Lesions in Rats and Mice |
| KO | Knock Out |
| LD | Lyme Disease |
| MC | Mast Cell |
| mL | Milliliters |
| PBS | Phosphate Buffered Saline |
| PFA | Paraformaldehyde |
| PRR | Pattern Recognition Receptor |
| PVI | Perivascular Infiltrate |
| $T_H2$ | Type 2 Helper T-cell |
| TLR | Toll-Like Receptor |
| μL | Microliters |
| US | United States |
| WT | Wild-type |

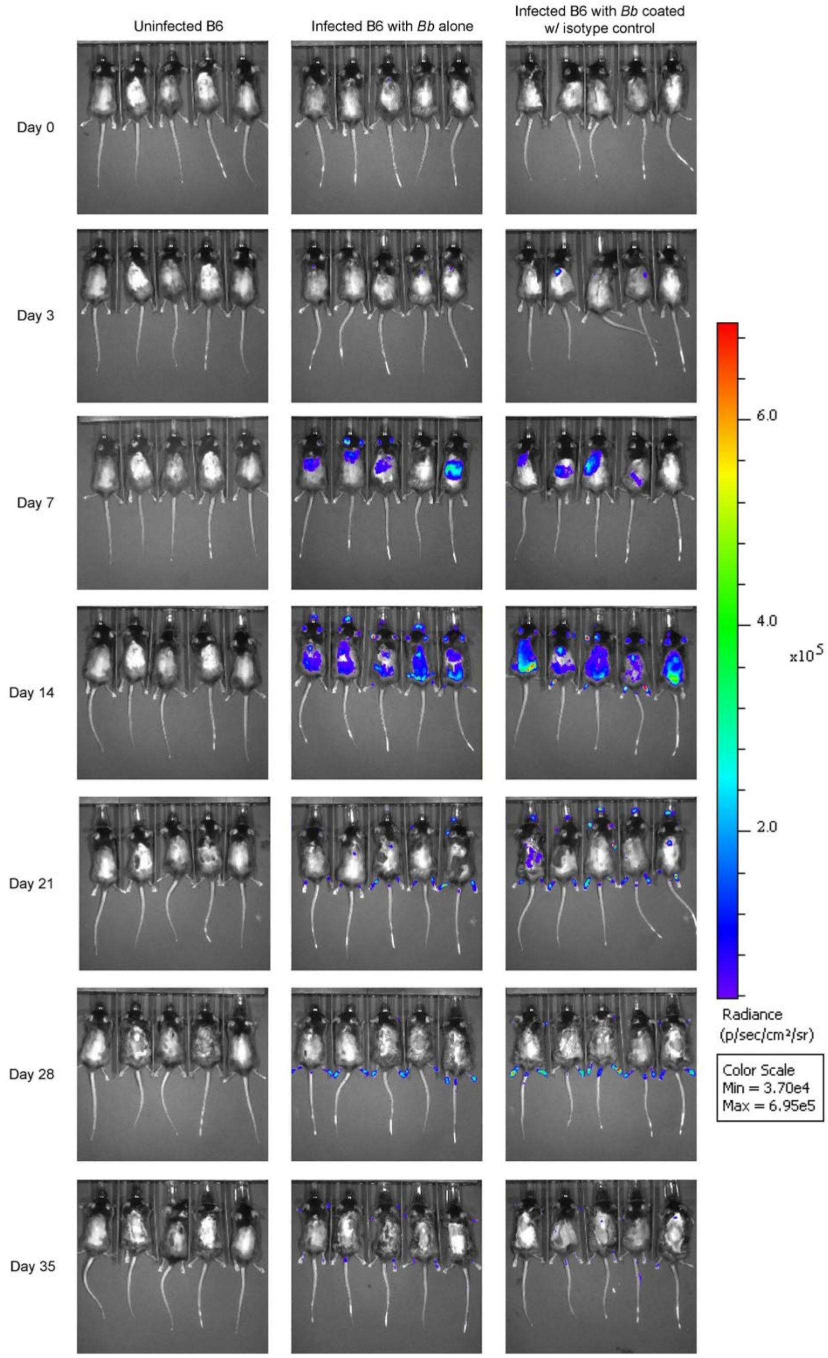


**Supplementary Figure 3. Bioluminescence image panel of C57BL/6J mice infected with ML23 *B. burgdorferi* in earlier time points.** Each mouse was infected with *B. burgdorferi* "ML23" containing the firefly *luc* gene encoding for luciferase. 100µL of D-luciferin was injected intraperitoneally in each mouse at 277mg/kg, then 15 minutes were allowed to pass as mice were shaved and Nair was applied to remove hair on the backs of the mice. Upon placement in the IVIS system, mice were imaged with an exposure time of 1 minute for bioluminescence.

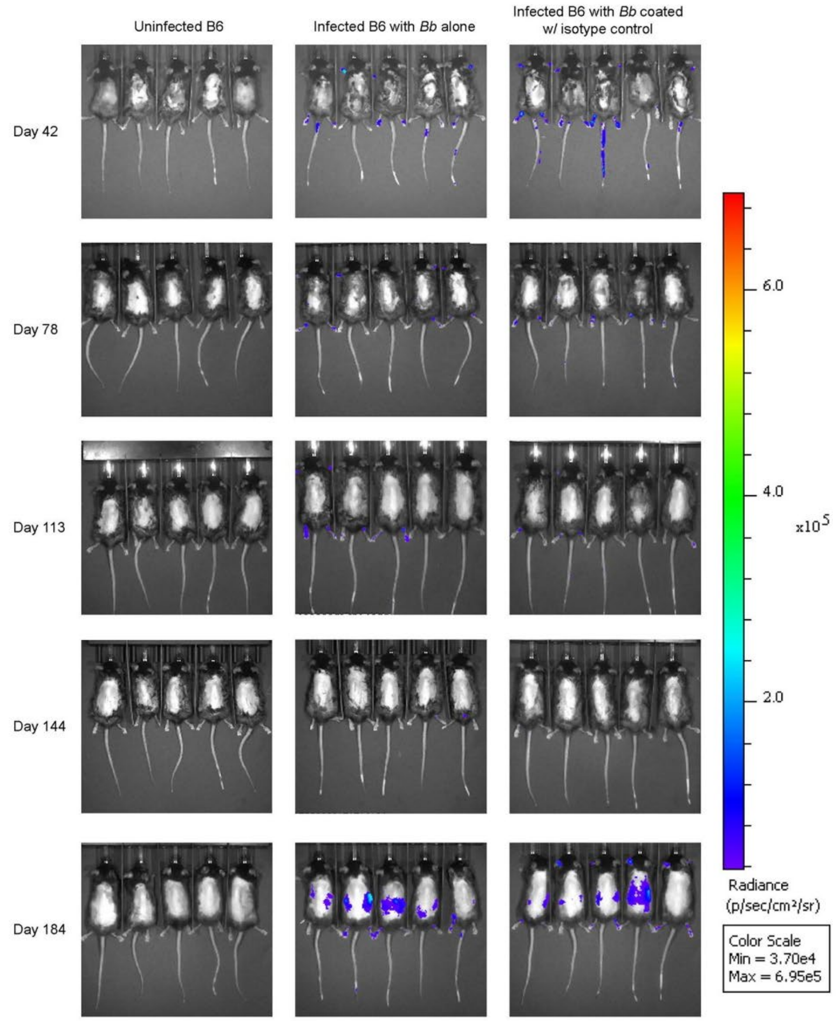


**Supplementary Figure 4. Bioluminescence image panel of C57BL/6J mice infected with ML23 *B. burgdorferi* in later time points.** Each mouse was infected with *B. burgdorferi* "ML23" containing the firefly *luc* gene encoding for luciferase. 100μL of D-luciferin was injected intraperitoneally in each mouse at 277mg/kg, then 15 minutes were allowed to pass as mice were shaved and Nair was applied to remove hair on the backs of the mice. Upon placement in the IVIS system, mice were imaged with an exposure time of 1 minute for bioluminescence.

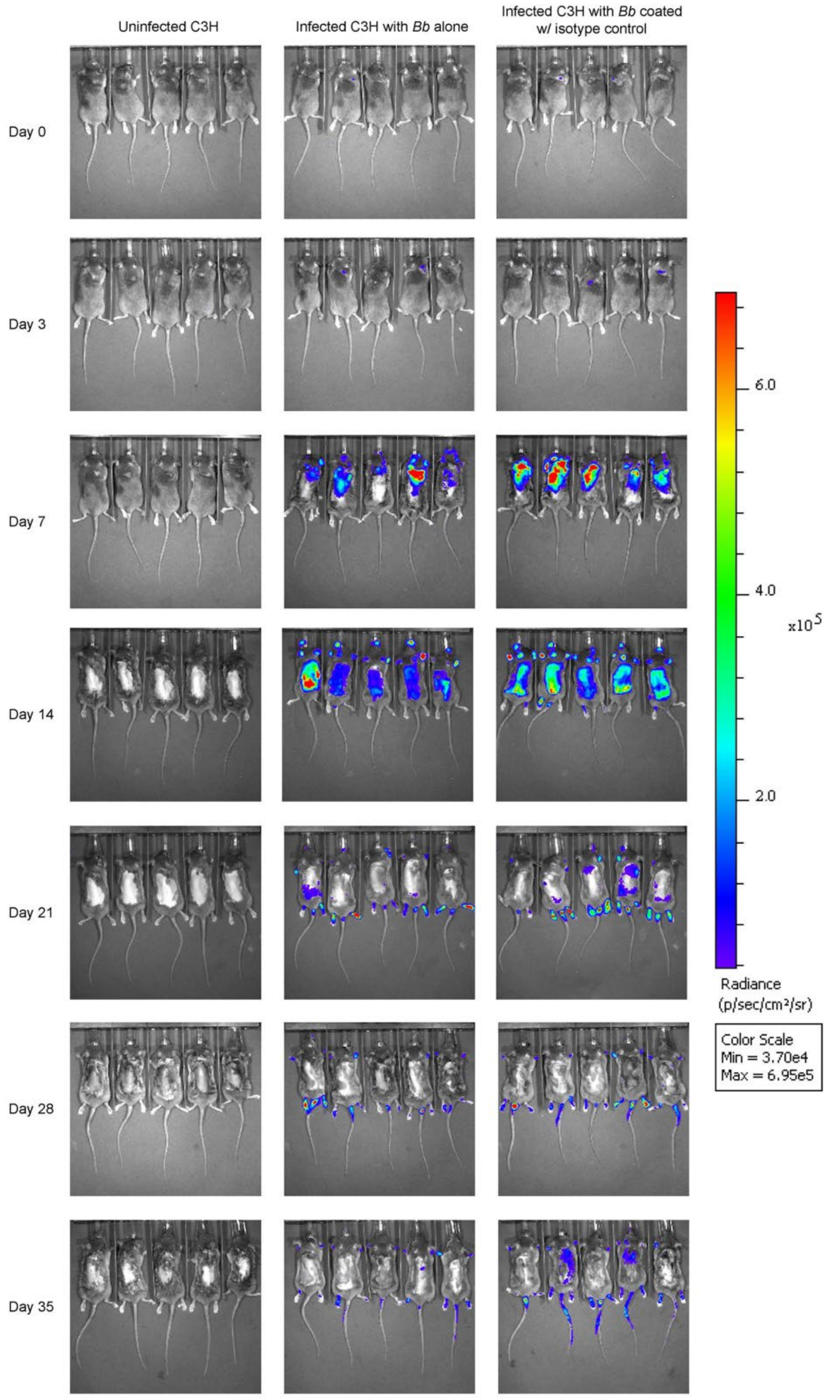


**Supplementary Figure 5. Bioluminescence image panel of C3H/HeJ mice infected with ML23 *B. burgdorferi* in earlier time points.** Each mouse was infected with *B. burgdorferi* "ML23" containing the fire-fly *luc* gene encoding for luciferase. 100µL of D-luciferin was injected intraperitoneally in each mouse at 277mg/kg, then 15 minutes were allowed to pass as mice were shaved and Nair was applied to remove hair on the backs of the mice. Upon placement in the IVIS system, mice were imaged with an exposure time of 1 minute for bioluminescence.

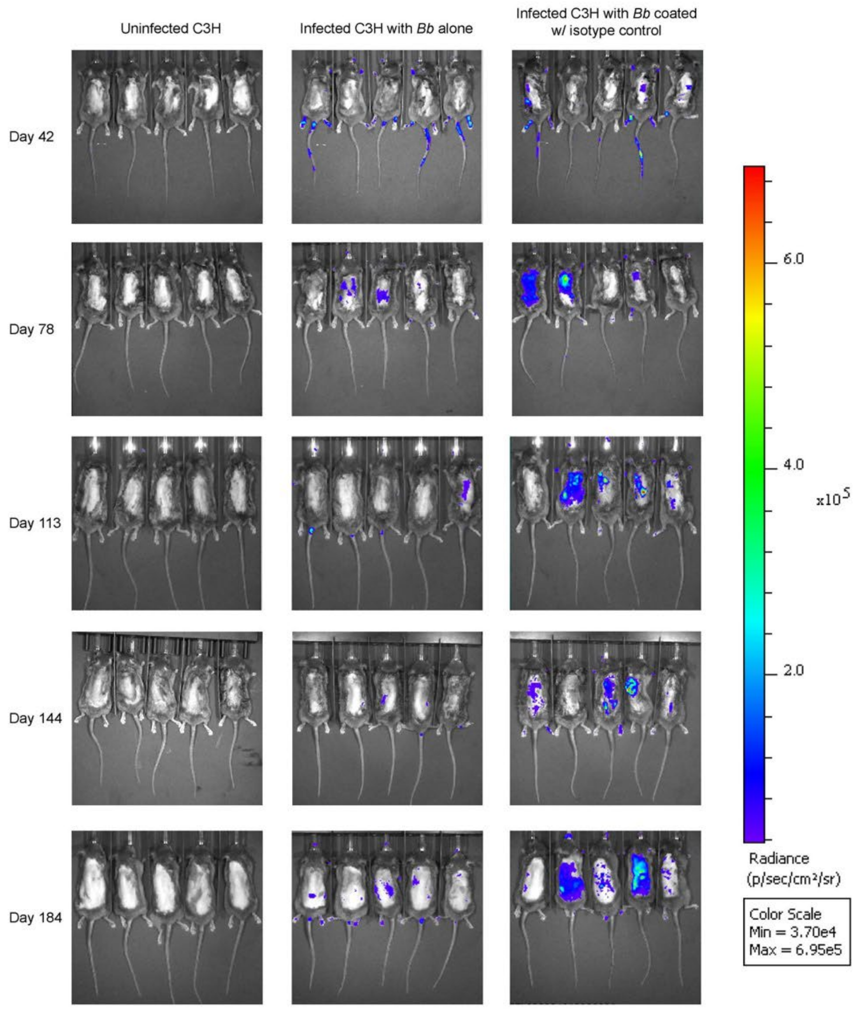


**Supplementary Figure 6. Bioluminescence image panel of C3H/HeJ mice infected with ML23 *B. burgdorferi* in later time points.** Each mouse was infected with *B. burgdorferi* "ML23" containing the firefly *luc* gene encoding for luciferase. 100µL of D-luciferin was injected intraperitoneally in each mouse at 277mg/kg, then 15 minutes were allowed to pass as mice were shaved and Nair was applied to remove hair on the backs of the mice. Upon placement in the IVIS system, mice were imaged with an exposure time of 1 minute for bioluminescence.
